# How flexible is tool use in Eurasian jays (*Garrulus glandarius*)?

**DOI:** 10.1101/803700

**Authors:** Piero Amodio, Markus Boeckle, Sarah A. Jelbert, Ljerka Ostoijc, Nicola S. Clayton

## Abstract

Eurasian jays (*Garrulus glandarius*) do not habitually use tools, yet they can be trained to solve object-dropping tasks, i.e. to insert a tool into an apparatus to release a food reward. Previous research suggests the these jays can learn a preference toward functional tools – objects allowing to obtain a food reward placed inside an apparatus – according to their density (Cheke et al., 2011). However, it is not yet known whether they can also select functional tools (tool selectivity) according to other physical properties such as size and shape, and use different kinds of tools to solve a similar task. Here we conducted three object-dropping experiments aimed at exploring these abilities in Eurasian jays. In Experiment 1, jays tended to select large stones as tools irrespective of the diameter of the apparatus. However, jays progressively developed a preference for the small tool, which was functional with both the wide and the narrow apparatuses. In Experiment 2, only vertically-oriented long stones could fit into the narrow apparatus, whereas both long and round stones were functional with the wide apparatus. Jays showed a preference for the long stone and, with the narrow apparatus, tended to achieve the correct manipulation after one or more unsuccessful attempts. In Experiment 3, jays were able to use sticks and adopt a novel technique on the same object-dropping apparatus, thus providing the first evidence that Eurasian jays can use sticks as tools. Taken together, these results indicate that Eurasian jays may have limited tool selectivity abilities but nonetheless can use different kinds of tools to solve similar tasks.

## Introduction

Corvids are a family of large brained birds thought to possess remarkable cognitive abilities (Emery and Clayton, 2004). One iconic expression of this sophisticated cognition is their skill in solving physical problems by using tools. The most prominent example is the New Caledonian crow (*Corvus moneduloides*), a species considered to be among the most proficient tool users in the animal kingdom. These birds are, together with Hawaiian crows (*Corvus hawaiiensis*), the only corvid species currently known to develop tool use behaviours in the absence of training (Kenward et al., 2005; Klump et al., 2018; Rutz, 2016). Individual practice and social inputs, however, appear to be essential for juvenile New Caledonian crows to acquire some of the more complex tool behaviours habitually performed in the wild, such as the manufacture of hooked stick tools or stepped pandanus tools (Holzhaider et al., 2010; Kenward et al., 2006). Growing evidence indicates that, although relying on an inborn predisposition for manipulating objects (for a review see Amodio et al., 2018), tool use behaviours in New Caledonian crows may entail complex physical cognition. For instance, tool manufacture in these crows varies across populations and represents an example of behavioural tradition (Hunt & Gray, 2003) that, according to a recent study, may be sustained through a mechanism of mental template matching (Jelbert et al., 2018). Furthermore, studies in captivity indicate that New Caledonian crows may be capable of flexibly selecting functional tools by encoding relevant features of objects: these crows have been reported to select or manufacture tools of appropriate length to acquire out-of-reach baits (Chappell and Kacelnik, 2002; Knaebe et al., 2017), choose the most suitable raw materials (e.g. plant species) to shape hook tools (Klump et al., 2019), and distinguish between light and heavy objects by observing the movement of objects in the breeze (Jelbert et al., 2019). New Caledonian crows have also been found to build composite tools (von Bayern et al., 2018), solve sequential tool use tasks (Taylor et al., 2007; Wimpenny et al., 2009), and – when presented with a multi-access apparatus – acquire food rewards by using up to four alternative strategies involving two different tools (i.e. sticks and balls) (Auersperg et al., 2011).

Corvid species not known to habitually use tools in the wild also exhibit impressive skills in solving problems by using tools. Possibly the most famous case are rooks (*Corvus frugilegus*). Bird and Emery (2009) conducted a series of object-dropping experiments demonstrating that these birds are capable of tool selectivity: they can select tools that are functional to solve a task, according to the physical properties of the tools (e.g. size, shape). Rooks were presented with a set of tools differing in one feature (e.g. size) and with an object dropping apparatus, a transparent box with a baited, collapsible platform in the inside and a vertical tube on the top. To solve the task, birds were required to select a functional tool (i.e. an object that could fit into the tube) and to drop it into the vertical tube of the apparatus, an action that would collapse the internal platform thus releasing the food reward. In the size selectivity test, rooks could choose between three stones of different sizes and they were tested in two conditions. In the first half of trials stones of all sizes were functional (Wide tube condition), whereas in the second half of trials only the small stones could fit into the apparatus (Narrow tube condition). Rooks were reported to have immediately switched their preference for large stones in the Wide tube condition to small stones in the Narrow tube condition. In the shape selectivity test, rooks were provided with two stones of different shapes and they were again tested in the Wide and Narrow tube conditions. Rooks selected and correctly oriented long stones (over non-functional round stones) in the Narrow tube condition, when only vertically-oriented long stones were functional (Bird and Emery, 2009). The authors found no evidence that rooks’ preference toward tools of appropriate size and shape emerged through learning, such that birds may have spontaneously adjusted their selection on the basis of the feature determining whether tools were functional or not. In a follow-up test, Bird and Emery (2009) showed that rooks can acquire food rewards from the same apparatus by using sticks, i.e. tools that differed from the ones they have been trained with, and that required a different technique. Rooks dropped heavy sticks in the same way they had previously dropped the stones, but they adopted a distinct technique to solve the task with light sticks: they held the tool with their beak and pushed it downward to collapse the baited platform. Through a further set of experiments, Bird and Emery (2009) found that rooks are also capable of solving a sequential tool use task, as well as fashioning functional tools by removing side branches from twigs or shaping hook-like tools from straight wire.

Common ravens (*Corvus corax*) are another example of corvids that can use tools in captivity, although they do not habitually exhibit such behaviours in the wild. Kabadayi and Osvath (2017) recently reported that one raven successfully solved an object-dropping task on the first training trial, and subsequently employed an alternative tactic when stones were unavailable: the bird filled the apparatus with parts of the aviary floor, and thereafter pecked at the substrate when it came within reach (Kabadayi and Osvath, 2017). Although anecdotal, this observation suggests that ravens, like New Caledonian crows and rooks, may be capable of using different kinds of tools and devising alternative strategies to acquire food from the same apparatus.

The flexibility in tool-use behaviours reported in these studies appears to indicate that complex physical cognition may be widespread within the *Corvus* genus, a subgroup of corvids to which rooks, common ravens, New Caledonian crows and Hawaiian crows belong. Why tool use in the wild is only exhibited by New Caledonian crows and Hawaiian crows may be explained by a number of factors, including stronger inborn predispositions for manipulating tools (Amodio et al., 2018; Kenward et al., 2005, 2006), as well as idiosyncratic features of their habitats (e.g. reduced risk of predation, lack of extractive foraging competitors; Rutz and St Clair, 2012).

To date however, it is not clear whether the sophisticated physical cognition reported in the *Corvus* genus is shared with more distantly related species of corvids. It has been shown that Eurasian jays (*Garrulus glandarius*) and California scrub-jays (*Aphelocoma californica*) can be trained to solve object-dropping tasks (Cheke et al., 2011; Logan et al., 2016; Miller et al., 2016). In the case of Eurasian jays, a member of the *Garrulus* genus, empirical evidence also suggests that these birds may take into account causal clues to solve tool use tasks. Cheke et al. (2011) presented Eurasian jays with a series of water displacement tasks that required the insertion of tools into an apparatus (i.e. a vertical tube filled with liquid) to acquire a reward, which could be reached only after the liquid has been progressively raised as a result of the insertion of the tools. In the study, the availability of causal cues was manipulated, such that in some tasks jays could choose between an apparatus or a tool that was functional according to physical principles and one that was not, whereas in other tasks jays had to select a functional apparatus according to arbitrary features such as colour, in the absence of available causal cues or when the available causal cues were counter-intuitive. Cheke et al. (2011) found that when causal cues were available, jays could learn to choose a functional liquid-filled apparatus over a non liquid-filled apparatus containing only air or filled with a solid substrate,, and functional sinking tools over ones that float and therefore fail to raise the water level. When the functional apparatus could be identified on the basis of arbitrary features rather than of causal cues, jays could also learn to select the functional apparatus but only in tasks in which the insertion of tools was ‘artificially’ (i.e. caused by the action of a hidden experimenter) associated with a progressive movement of the food reward. In contrast jays failed to learn a preference toward the functional apparatus in a counter-intuitive task. When presented with a modified apparatus formed by one baited narrow tube and two non-baited wide tubes with different colour marks, jays could not learn to drop tools into the functional non-baited tube, an action that would raise the level of the liquid not only in the functional non-baited tube, but also in the adjacent baited tube because these tubes were invisibly connected. Thus, the overall performance of Eurasian jays in this study suggests that these corvids may have acquired tool use behaviours through an interplay between instrumental learning and causal understanding, with the latter fostering/constraining the learning process according to whether tasks fit or do not fit with physical principles (Cheke et al., 2011).

It is important to note is that in the Cheke at al. (2011) study there are two confounding variables which may have affected jays’ performance in the task requiring jays to select functional sinking objects over non-functional floating objects. First, the two kinds of objects differed not only in density, the physical property which determined its functionality, but also in material (rubber or foam). Second, both individual birds that were tested had previous experience in dropping rubber tools, but not foam tools, in order to solve water displacement tasks. Specifically, both birds had taken part in a previous water displacement experiment in which rubbers and stones were provided as tools. Note however that their amount experience was quite different: one bird, Hoy, had used rubber on 35 occasions, whereas the other bird, Romero, had used rubber only once, and yet Hoy’s performance was no better than Romero’s. Nonetheless, it cannot be excluded that jays developed a preference toward the functional rubber tool on the basis of trivial features (e.g. material) or due to a higher familiarity with these objects. Thus it remains an open question whether Eurasian jays are capable – as rooks and New Caledonian crows – of selecting tools on the basis of an understanding of functionality.

To overcome these potential confounds and limitations, here we conducted two tool selectivity tests in which functional and non-functional tools differed only in one feature (i.e. size in Experiment 1, and shape in Experiment 2), and were not familiar to the birds in the context of object dropping tasks. We presented the birds with two object dropping tasks that were designed after Bird and Emery’s (2009) size and shape selectivity tests, and which involved an apparatus closely resembling that used in rooks. Finally in a third experiment we explored jays’ capability of using different tools – sticks – to acquire food for the same apparatus.

## METHODS

### Subjects

Five hand-raised Eurasian jays of both sexes were tested (Chinook, Homer, Jaylo, Poe, Stuka). At the time of testing (October-December 2016), all birds were juveniles (1.5 years). Birds were group-housed in a large outdoor aviary (20×10×3m) at the Sub-Department of Animal Behaviour, University of Cambridge. The birds received a maintenance diet of vegetables, eggs, seeds and fruits and water *ad libitum*. All birds took part in Experiment 1a. Chinook stopped interacting with the apparatus and the tools after completing this experiment, therefore she was excluded from subsequent experiments. All birds except Chinook were tested in the subsequent Experiments 1b, 2 and 3.

### Apparatus

All tests were conducted using an object dropping apparatus originally designed by Bird & Emery (2009) and modified for Eurasian jays by Cheke et al. (2011) and Miller et al. (2016). It consisted of a transparent Perspex box (12×11×11 cm) with a baited but out-of-reach, collapsible platform with a vertical tube (11.5 cm) on top (Fig. 1). Depending on the experimental condition, either a wide tube (Ø 4.2 cm) or a narrow tube (Ø 1.6 cm) was used. To release the food, birds could drop a tool (e.g. a stone) into the tube, which then caused the internal platform to collapse and food to fall out.

**Figure 1:**
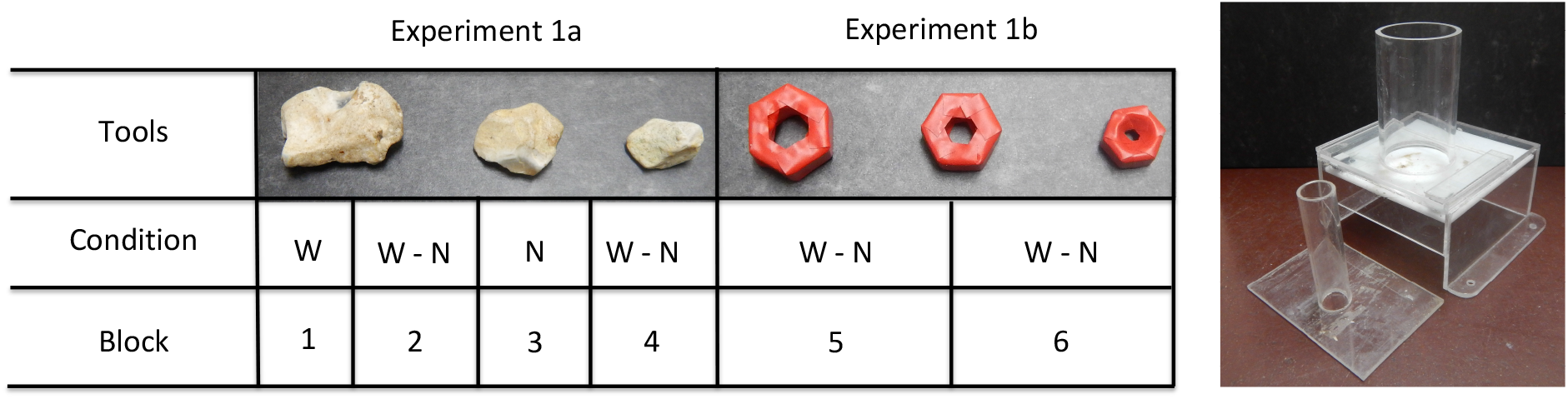
Design and Apparatus used in Experiment 1. Left) Scheme showing the tools (stones or novel objects) and conditions (Wide tube, W; Narrow tube, N) used in each block (20 consecutive trials) in Experiment 1a, and 1b; Right) Picture of the Wide tube apparatus (back) and Narrow tube (front).

### General procedure

Birds were tested in visual isolation from other individuals inside an indoor compartment (2×1×3 m). The experimenter placed the apparatus and the tools into the bird’s compartment through an opening in the mesh wall. In all three experiments, the tools were placed on one side of the apparatus, approximately 10 cm away from it. The initial position of the bird was not standardized, so that the bird could be in any location within the compartment at the onset of each trial, i.e. when the stimuli were presented. In experiments involving multiple tools (Experiments 1 and 2), the tools were equidistant from the bird when the bird was facing the front side of the apparatus (i.e. the side of the apparatus from which the food is released). In Experiments 1 and 2, the position of the tools in regards to the apparatus (i.e. close, middle, far position) was pseudo-randomized across trials. Live larvae of the mealworms beetle (*Tenebrio molitor*) were used as food rewards. The baiting of the apparatus occurred out of view.

In all experiments the maximum duration of a trial was set to 2 minutes. However, in Experiment 3, one additional minute was allowed if the bird was interacting with the tool by the cut-off time (up to 4 minutes in total). This procedural detail was set during the very first trial, when it was noticed that the jay being tested required substantially more efforts to achieve the correct manipulation of stick tools.

Typically, birds were given 10 trials per day. However, if a bird did not interact with the tool(s) at all during a trial, testing was interrupted and continued on the next day, with the no-interaction trial being repeated. For example if a bird completed trials 1-8 and subsequently it stopped interacting with the stimuli on trial 9, then on the day the bird received trial 9 again, and potentially 10 additional trials (i.e. trials 9-18).

Subjects did not have access to their maintenance diet for 1h prior to testing to ensure that they were motivated to eat multiple food rewards during testing. Water was accessible *ad libitum* during testing. All experiments were recorded using a GoPro^®^ Hero 4 video-camera and subsequently analysed.

### Specific Procedure and Materials

#### Refresh Training

All birds had previously been trained to drop hollow metal balls (Ø 2 cm, 4 g) inside the apparatus as part of a previous study (Miller et al., 2016). All birds had also been further exposed to the wide tube apparatus and metal balls in a non-systematic manner. This occurred during January-April 2016, as part of the training of one of us (PA) in working with the jays. Critically, however, the birds had no prior experience of dropping any of the specific tools used in this study (stones and sticks).

We conducted a short refresher training to ensure that the birds were still familiar with the task and would insert stones into the apparatus. During training, the birds were presented with the wide tube apparatus and a single tool placed approximately 10 cm away from it. In the first 5 trials, they were provided with the metal ball (i.e. the tool with which they had previously been trained) and in the following 5 trials with a medium stone (11.2 ± 0.4 g). All birds successfully solved these 10 trials before testing started.

### Experiment 1: Size Selectivity Test

The design of this experiment closely followed the ‘Stone Size Test’ conducted by Bird and Emery (2009) on rooks. We used both the wide and the narrow tube apparatuses in the test. In Experiment 1a, stones of three sizes – large (7.8 ± 0.2 g), medium (4.2 ± 0.5 g), small (2.2 ± 0.1 g) – were provided as tools. With the wide tube, stones of all sizes were functional (i.e. all stones could fit inside the tube), whereas with the narrow tube, only the small stone was functional because the two larger stones did not fit inside the tube. The experiment was composed of four blocks (Fig. 1). In Block 1 (trials 1-20), jays were presented only with the wide tube apparatus to evaluate their potential preference for a specific tool, namely the small versus the large stone. In Block 2 (trials 21-40), the narrow tube (10 trials in total) and the wide tube (10 trials in total) were presented in a pseudo-randomized order to investigate whether the jays would spontaneously select the small stone when this was the only functional tool (Narrow tube condition), and if they would express the same preference when all stones were functional (Wide tube condition). In Block 3 (trials 41-60) only the narrow tube apparatus was used. This block was designed to facilitate jays’ learning about the functional features of the small stone. Finally, in Block 4 (trials 61-80) birds received a further test with the narrow and wide tubes that followed the procedure of Block 2. This experiment differed from the ‘Stone Selectivity Test’ conducted by Bird and Emery (2009) in two respects. Rooks tested in the latter study received a smaller number of trials (i.e. 60 trials), and they experienced only the wide tube apparatus in the first 30 trials and only the narrow tube apparatus in the remaining 30 trials.

After noting that the jays appeared to have switched to using the small stone more often in Block 4 than in Block 2, in Experiment 1b we investigated whether jays may have learned to select the small stone based on its functional property, namely its size. To that end, Experiment 1b was a transfer task with novel objects, which differed from the stones in irrelevant perceptual properties namely colour, shape and material. In the same way that the stones previously differed in their functional property of size, the novel objects were large (15 g), medium (10.5 g) and small (5 g) bolts upholstered with red tape. Like previously, the novel objects of all sizes were functional with the wide tube but only the small one could fit in the narrow tube. Given that jays had had two blocks of the two apparatuses in a counterbalanced order in Experiment 1a, they also received two blocks in Experiment 1b (Fig. 1): Block 5 (trials 81-100), Block 6 (trials 101-120). In each blocks, there were 10 trials with the narrow tube in total and 10 trials with the wide tube, the order of which was pseudo-randomized such that the jays did not receive more than three consecutive trials with the wide tube nor narrow tube.

### Experiment 2: Shape Selectivity Test

The design of this experiment closely followed the ‘Stone Orientation Test’ used by Bird and Emery (2009) for rooks. In Experiment 2, jays were provided with two shapes of stone tools: long stones (approximately 2.4×1.0 cm) and round stones (approximately 1.8×1.9 cm). Birds received a total of 20 trials, 10 of which were with the narrow tube and 10 of which were with the wide tube. The order in which birds were presented with the narrow and wide tubes was pseudo-randomized such that the jays did not receive more than three consecutive trials with the wide tube nor narrow tube. To successfully solve the task in the Narrow tube condition, birds had to select the long stone and orient it vertically to insert it into the tube. In the wide tube condition, both stones were functional and no specific rotation of the tool was required. This experiment and the equivalent test previously conducted on rooks (Bird and Emery, 2009) differed in the number trials (40 trials in rooks) and in the fact that the two apparatuses were not counterbalanced within sessions of trials (rooks first received 20 trials with the wide tube apparatus, and subsequently 20 trials with the narrow tube apparatus).

### Experiment 3: Stick Tool Test

The design of this experiment closely matched the ‘Stick Use Test’ conducted by Bird and Emery (2009). In Experiment 3, jays were provided with one of two types of sticks as a tool: a twig (11 cm long, 3.0 g) or a barbecue stick (11 cm long, 0.4 g). When provided with the twig, jays could solve the task by dropping it, just like they previously did with stones (*No Push* technique). Due to its lighter weight, the barbecue stick required jays to hold it in their beak and push it downwards to collapse the baited platform (*Push* technique).

A total of 10 trials with the wide tube were conducted. On each trial, jays were only provided with one type of tool (the barbecue stick or the twig), the order of which was pseudo-randomized such that jays did not receive the same tool on more than two consecutive trials. In contrast to this experiment, rooks tested by Bird and Emery (2009) received more trials (20 trials) and they were presented consistently with the heavy stick in the first 10 trials, and then the light sticks in the final 10 trials.

### Data Analysis

All data were analysed with R.3.5. using the RStudio 1.1.447 wrapper (RStudio Team, 2018). In Experiment 1 we scored the tool (small, medium or large) selected on each trial. Ordinal-logit models (package *ordinal*, Haubo and Christensen, 2018) were used to test whether jays adjusted their preference of selection (i) according to the condition (Wide or Narrow tube), and (ii) across blocks (e.g. due to learning). In all models, the size of the tool (small, medium or large) selected in each trial was treated as ordinal data and fitted as response variable.

In Experiment 2 we scored the tool (long or round stone) selected and what kind of manipulation of the long stone was performed on each trial. Specifically, three kinds of manipulation could be achieved. The long stone could: i) be oriented vertically prior of the first insertion attempt (*Immediate Rotation*); ii) be oriented vertically after one or more failed insertion attempts (*Eventual Rotation*), or; iii) not be oriented vertically (*No Rotation*). The scoring of these behaviours had been planned before the experiment was conducted, based on the results previously reported in rooks on this test (Bird and Emery, 2009). Binomial General Linear Models (package *stats*, R Core Team and contributors worldwide, 2018) were fitted to test whether the kind of tool selected and the manipulation performed varied (i) according to the condition and, (ii) across blocks (e.g. due to learning).

In Experiment 3 we scored: i) whether a trial was successful, ii) the technique utilized to solve the task with stick tools (i.e. *Push* technique, *No Push* technique), and iii) the number of insertion attempts until successfully inserting the tool into the apparatus. The scoring of successful trials and the tool use technique had been planned before the experiment was conducted, based on the results previously reported in rooks on this test (Bird and Emery, 2009). We decided to score the number of insertion attempts during the testing phase as it became clear that this variable was very informative of inter-individual differences in performances. The data were analysed descriptively.

All datasets and R scripts used to conduct the statistical analyses are available on Zenodo (https://doi.org/10.5281/zenodo.3471706)

## RESULTS

### Inter-observer Reliability

PA coded all videos and Benjamin Farrar (BF) acted as naïve coder, analysing 20% of videos for each experiment. Inter-observer reliability was excellent: for Experiment 1 (size of selected tools: Cohen’s kappa *k*=0.986, p <0.0001) and Experiment 2 (shape of selected tools: Cohen’s kappa *k*=1, p <0.0001; kind of rotation of the tool: Cohen’s kappa *k*=1, p <0.0001), and for Experiment 3, PA and BF achieved 100% of agreement in scoring successful trials and insertion techniques.

### Experiment 1: Size Selectivity Test

In Block 1, one bird (Stuka) received 15 instead of 20 trials due to an experimental error. Overall jays selected a similar proportion of the three stones between conditions (Model 1, Tab. 1). Hence jays did not adjust the selection of tools according to the diameter of the tube. We further tested whether the proportion of selection of the three stones varied across blocks by comparing jays’ performance in Block 1 with each of the three subsequent blocks. The performance in Block 1 represents the spontaneous preference exhibited by the jays when all tools were functional, therefore this block was considered as a meaningful reference to analyse changes of preference through dyadic comparisons among blocks. Model 1 (Tab. 1) showed that the proportions of selection of the three stones in Block 1 were comparable to those observed in the two subsequent blocks, but significantly different from the proportions of selection in Block 4. This result is likely to be explained by the variation in preference for the large and the small stones throughout blocks (Fig. 2). A stronger preference towards the large stone appeared in Block 1 and was retained across Block 2 and 3, but it became less pronounced together with an increase in preference for the small stone in Block 4 (Fig. 2).

**Figure 2:**
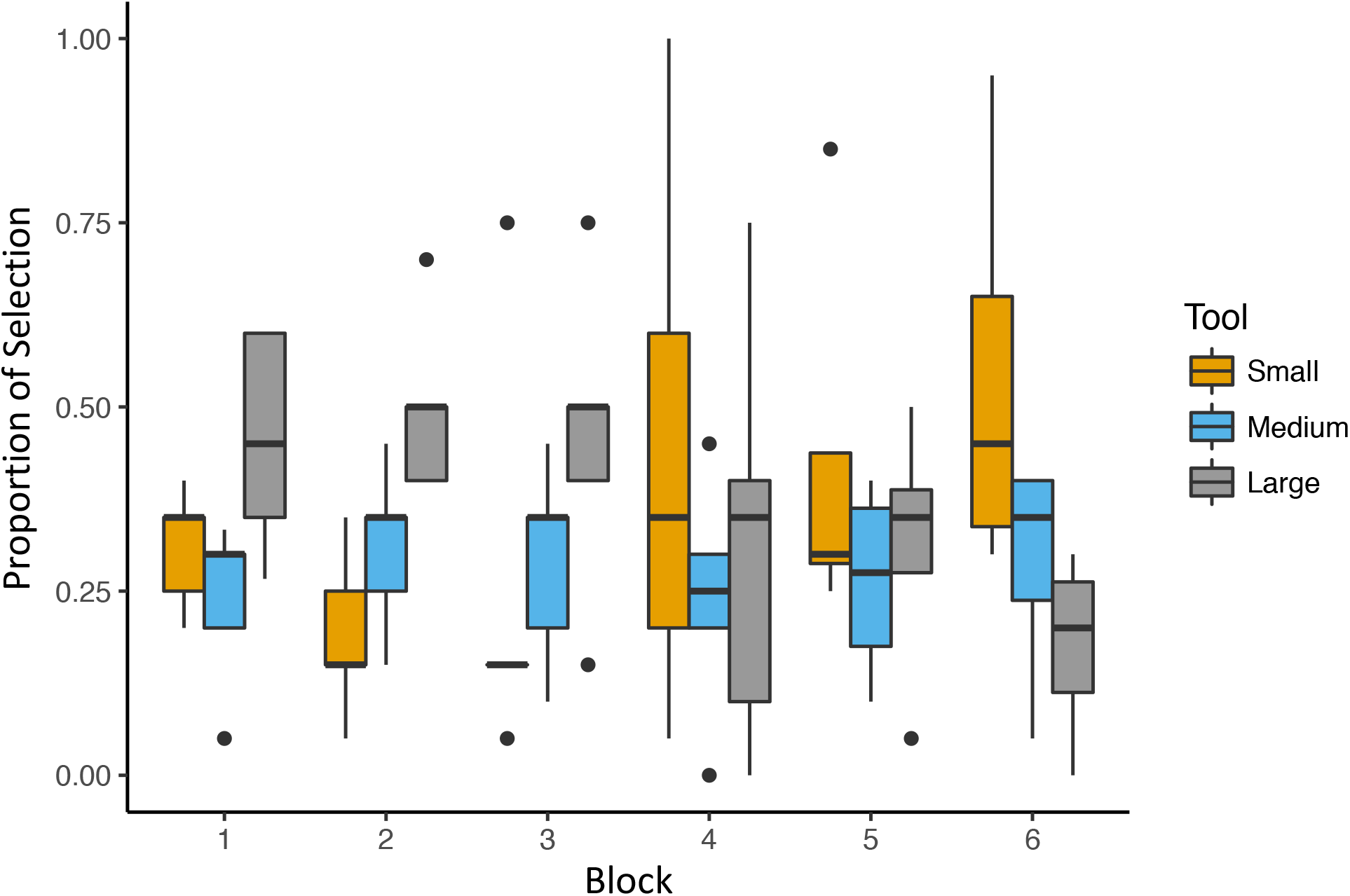
Box and Whisker plot showing the proportions of selection of the small, medium and large tool in Experiment 1a (Block 1-4) and Experiment 1b (Block 5,6). In Blocks 2, 4, 5, and 6, the two conditions (Narrow and Wide tube) are grouped as jays’ performance was comparable across conditions (Model 1, Tab. 1).

**Table 1:**
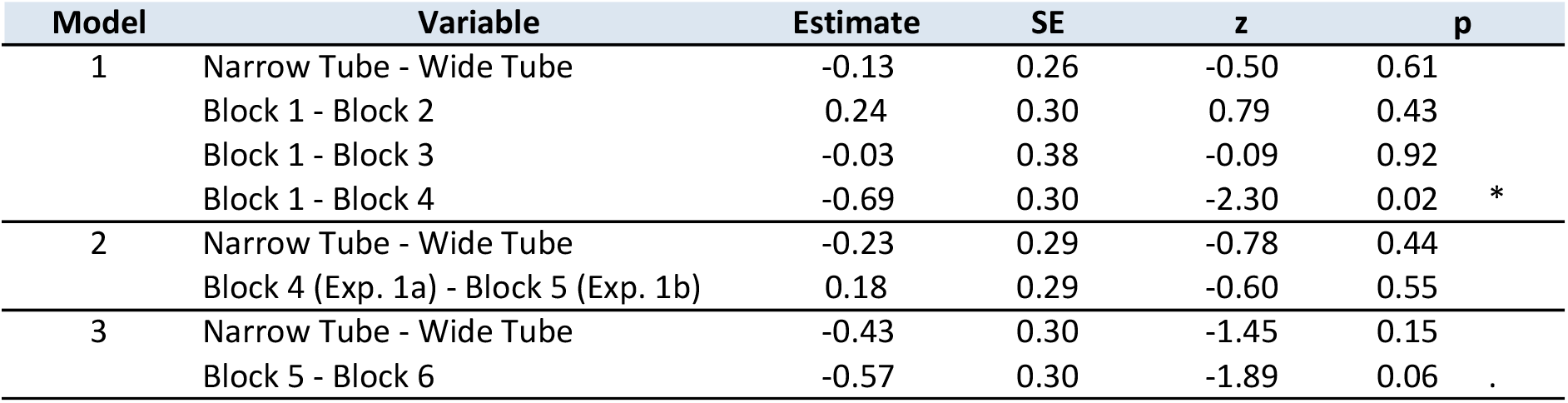
Results from ordinal-logit models examining whether the proportions of selection of the three tools differed: i) according to the condition (Wide or Narrow tube), and; ii) across blocks of trials. Model 1 focused on Experiment 1a. Model 2 focused on the last block of Experiment 1a and the first block of Experiment 1b. Model 3 focused on Experiment 1b. SE: standard error; z : z value; p: p value.

| Model | Variable | Estimate | SE | z | p |
| --- | --- | --- | --- | --- | --- |
| 1 | Narrow Tube - Wide Tube | -0.13 | 0.26 | -0.50 | 0.61 |
|  | Block 1 - Block 2 | 0.24 | 0.30 | 0.79 | 0.43 |
|  | Block 1 - Block 3 | -0.03 | 0.38 | -0.09 | 0.92 |
|  | Block 1 - Block 4 | -0.69 | 0.30 | -2.30 | 0.02 * |
| 2 | Narrow Tube - Wide Tube | -0.23 | 0.29 | -0.78 | 0.44 |
|  | Block 4 (Exp. 1a) - Block 5 (Exp. 1b) | 0.18 | 0.29 | -0.60 | 0.55 |
| 3 | Narrow Tube - Wide Tube | -0.43 | 0.30 | -1.45 | 0.15 |
|  | Block 5 - Block 6 | -0.57 | 0.30 | -1.89 | 0.06 . |

To investigate whether the slight shift in the jays’ preference for the small tool may have been due to its functional feature represented by size, we compared the jays’ performance in the last block when tools were stones (Experiment 1a; Block 4) with the first block when tools were novel objects (Experiment 1b: Block 5). One subject (Chinook) was excluded from this analysis because she did not participate in Experiment 1b. Consistent with previous analysis, jays’ performance was comparable across conditions (Model 2, Tab. 1). Model 2 also indicated that the proportions of selection of the three stones in Block 4 were similar to the proportions of selection of the three novel objects in Block 5 (Tab. 1; Fig. 2). This means that jays showed a similar pattern of behaviours with the stones and the novel objects.

Together, these results are consistent with the possibility that jays transferred their pattern of preference between tools of different appearances. However, because we did not test jays’ preference for the three novel objects in a naïve group of birds, we cannot rule out the possibility that jays’ performance in Block 5 resulted from a difference in spontaneous preference for the novel objects over stones that was not influenced by their experience in Experiment 1a.

Finally, we compared jays’ preference of selection of the three novel objects between the two blocks of Experiment 1b. Model 3 (Tab. 1) revealed that the proportions of selection of the three tools in Block 5 and Block 6 were not significantly different, thus indicating that jays’ preferences of selection was stable across blocks. However, the model showed a trend (p< 0.06), which could be explained by the concurrent stronger preference for the small tool and decreased preference for the large tool in Block 6 (Fig. 2). This pattern first appeared in Block 4, was then retained in Block 5, and finally increased – although not significantly – in Block 6 (Fig. 2). Supporting previous findings, Model 3 also indicated that jays’ performance was again consistent between conditions.

Diagnostic plots were produced for each of the three models. Visual inspection of the plots indicates that all models have a satisfactory fit.

### Experiment 2: Shape Selectivity Test

When presented with a choice of two stones of different shapes, jays showed a pronounced preference for the long stone in both conditions (Narrow tube: 77.5 ±7.5% trials; Wide tube: 77.5 ±4.8% trials; Mean ± SE). The GLM analysis indicated that jays’ preference for the long stone was stable across conditions and blocks (Model 4, Tab. 2).

**Table 2:** Results from GLMs examining data of Experiment 2. Model 4 tested whether the proportions of selection of the tools (long and round stones) differed: i) according to the condition (Wide or Narrow tube), and; ii) across blocks of trials. Model 5 tested whether the proportions in which the long stone was oriented vertically (i.e. *Immediate Rotation*, *Eventual Rotation*) or it was not oriented vertically (*No Rotation*) differed: i) according to the condition (Wide or Narrow tube), and; ii) across blocks of trials.

| Model | Variable | Estimate | SE | z | p |  |
| --- | --- | --- | --- | --- | --- | --- |
| 4 | Intercept | -1.38 | 0.47 | -2.94 | 0.003 | ** |
|  | Narrow Tube - Wide Tube | -0.01 | 0.54 | -0.03 | 0.98 |  |
|  | Block 1 - Block 2 | 0.29 | 0.54 | 0.53 | 0.60 |  |
| 5 | Intercept | 1.50 | 0.53 | 2.81 | 0.005 | ** |
|  | Narrow Tube - Wide Tube | -2.16 | 0.59 | -3.63 | 0.0003 | *** |
|  | Block 1 - Block 2 | 0.16 | 0.59 | -0.27 | 0.79 |  |

In line with previous findings in rooks (Bird and Emery, 2009), the jays performed three kinds of manipulation when the Long stone was selected. The tool was oriented vertically (Fig. 3) either prior to the first insertion attempt (*Immediate Rotation*) or after one or more unsuccessful attempts (*Eventual Rotation*). Alternatively the tool was oriented horizontally with respect to the tube (*No Rotation*, Fig. 3).

**Figure 3.**
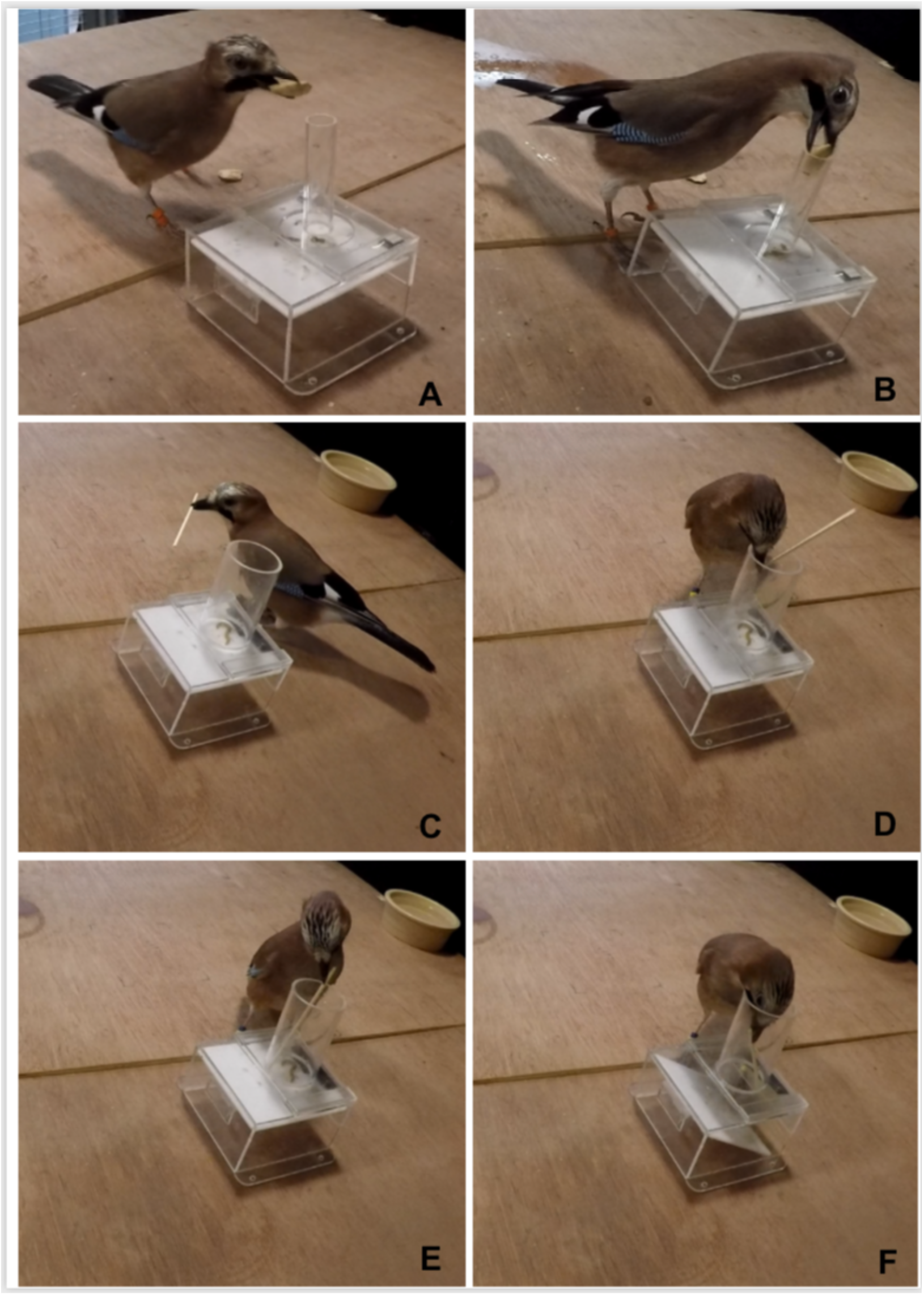
Eurasian jay being tested in Experiment 2 and Experiment 3: A-B) Different kinds of manipulation of the Long stone: *No Rotation* (A), Rotation (B); C-F) Sequence of actions describing the *Push* technique: the stick is picked up near one end (C), steered inside the tube (D) and pushed downward (E) causing the collapse of the internal platform of the apparatus (F).

To investigate whether the manipulation of the long stone differed between conditions we fitted a GLM with a binary outcome variable (*No Rotation*-Rotation). The manipulation of the long stone differed between the conditions (Model 5, Tab. 2) with higher frequencies of rotation performed with the Narrow tube (Narrow tube: 79.1 ±10.1% trials; Mean ± SE). However, the correct orientation of the long stone in the Narrow tube condition was often achieved *after* one or more incorrect attempts of insertion (*Eventual Rotation*: 55.3 ± 21.1% of trials with the narrow tube in which the long stone was rotated; Fig. 5). Therefore this finding cannot be taken as evidence that jays had a solid understanding of the affordance of the task, because in this case they would have correctly oriented the stone before the first insertion attempt (*Immediate Rotation*). Model 5 (Tab. 2) also indicated that the manipulation of the long stone was stable across blocks.

### Experiment 3: Stick Tool Test

All subjects solved the task by using a stick as a tool (Tab. 3). However, the insertion rate was extremely variable: Homer 9/10 trials, Jaylo 8/10 trials, Poe 2/10 trials, Stuka 1/10 trials. Similarly, the number of insertion attempts was also quite variable among subjects (Tab. 3). The success rate matched the insertion rate for all subjects except for Jaylo, who did not collapse the platform in 3 trials in which the tool was inserted into the apparatus.

**Table 3:** Trial by trial description of the behaviour in Experiment 3. In successful trials (grey cells), the technique that was used is reported. Blank cells correspond to unsuccessful trials.

| Trial | Homer |  |  |  | Jaylo |  |  |  | Poe |  |  |  | Stuka |  |  |  |
| --- | --- | --- | --- | --- | --- | --- | --- | --- | --- | --- | --- | --- | --- | --- | --- | --- |
|  | Stick Kind | Inserted | N°Attempts | Technique | Stick Kind | Inserted | N°Attempts | Technique | Stick Kind | Inserted | N°Attempts | Technique | Stick Kind | Inserted | N°Attempts | Technique |
| 1 | BBQ Stick | Y | 2 | Push | BBQ Stick | N | 14 |  | BBQ Stick | N | 16 |  | Twig | N | 1 |  |
| 2 | Twig | N | 5 |  | Twig | Y | 12 | Push | Twig | Y | 11 | No Push | Twig | Y | 5 | Push |
| 3 | BBQ Stick | Y | 2 | Push | BBQ Stick | N | 12 |  | BBQ Stick | N | 16 |  | BBQ Stick | N | 4 |  |
| 4 | BBQ Stick | Y | 2 | Push | BBQ Stick | Y | 7 |  | BBQ Stick | Y | 9 | Push | Twig | N | 4 |  |
| 5 | Twig | Y | 2 | No Push | Twig | Y | 7 | Push | Twig | N | 15 |  | Twig | N | 2 |  |
| 6 | Twig | Y | 13 | Push | Twig | Y | 3 | Push | Twig | N | 0 |  | BBQ Stick | N | 0 |  |
| 7 | BBQ Stick | Y | 1 | Push | BBQ Stick | Y | 9 |  | BBQ Stick | N | 2 |  | BBQ Stick | N | 0 |  |
| 8 | Twig | Y | 3 | Push | Twig | Y | 5 | Push | Twig | N | 0 |  | Twig | N | 0 |  |
| 9 | Twig | Y | 1 | No Push | BBQ Stick | Y | 8 |  | BBQ Stick | N | 0 |  | BBQ Stick | N | 0 |  |
| 10 | BBQ Stick | Y | 3 | Push | Twig | Y | 1 | Push | Twig | N | 0 |  | BBQ Stick | N | 0 |  |

In most of the successful trials (77%), jays collapsed the platform of the apparatus by actively pushing downward on the stick (*Push* technique). The technique was adopted not only with light barbecue sticks but also with twigs. In 23% of successful trials, the insertion of the twig into the apparatus was sufficient to collapse the platform in the absence of active pushing (*No Push* Technique). The reason for the active pushing with the heavy stick can likely be explained by the insertion technique used by the birds. Instead of dropping the stick as they did with stones, birds typically held the stick near one end and appeared to carefully steer it inside the tube (Fig. 3). As a result of these seemingly gentle movements, the heavy stick likely did not always hit the platform with enough force to collapse it. All birds used the *Push* technique at least once. Two subjects (Homer and Poe) solved the task by using both techniques.

## DISCUSSION

Here we investigated the tool use abilities of Eurasian jays by exploring whether this species of corvid can select appropriate tools on the basis of their physical properties, namely size and shape, and solve a familiar task by using novel tools, sticks. Jays showed only limited tool selectivity, i.e. they did not spontaneously adjust their choice according to the functionality of the tools, but were capable of using sticks as tools.

In the size selectivity test (Experiment 1), jays initially exhibited a spontaneous preference for the large stone regardless of whether this tool was functional or not. Thus, jays seem to have failed to encode the relevant feature of objects and to adjust their selection of tool according to the features of the apparatus. However, jays’ performance also suggests that they may be capable of altering their selection of tools through learning: across trials jays reduced their initial preference in favour of the only tool that was functional in both conditions, namely the small stone. Subsequently, when presented with novel objects (Experiment 1b), jays expressed a pattern of selection that was comparable to those observed toward stones in the final block of the previous test (Experiment 1a). This result is consistent with the possibility that jays transferred their preference of which tool to use based on size despite other perceptual differences between the objects (perhaps through a process of generalization, cf. Shettleworth, 2010).

In the shape selectivity test (Experiment 2) jays exhibited a pronounced preference for the functional tool (long stone) in both conditions, but they tended to perform the correct manipulation of the tool only when needed (Narrow tube condition). However, jays often achieved the correct manipulation of the tool after one or more failed attempts of insertion (*Eventual Rotation*) rather than before the first insertion attempt (*Immediate Rotation*). Therefore, it is likely that jays correctly oriented the tools through trial-and-error, in the lack of a full understanding of the objects’ properties and functionality.

In the stick tool test (Experiment 3), all birds were capable of using a novel tool and of acquiring food rewards from a familiar apparatus through a novel strategy (i.e. *Push* technique). These results also represent the first demonstration that Eurasian jays can use sticks as tools.

The overall pattern of our results support previous reports of learning forming the basis of Eurasian jay tool use (Cheke et al. 2011). The relatively fast learning of a preference for the functional tools in the latter study may have been facilitates by the fact that birds had already experienced the functional sinking objects as tools, before the study (Cheke et al. 2011) and that the functional and non-functional tools differed not only in the relevant characteristic (density) but also in others features (material and colour).

Further in line with Cheke et al. (2011), we found important individual differences in tool use skills in Eurasian jays. One bird, Homer, rapidly developed a clear preference for the functional small tool in Experiment 1 (Fig. 4) compared with the other jays that were tested; Homer more frequently oriented the tool correctly *before* the first insertion attempt (*Immediate Rotation*) in Experiment 2 (Fig. 5) and solved more trials (9/10) and by using both techniques in Experiment 3 (Tab. 3). A possible explanation for the performance of this individual may be linked to his experimental history. Homer was used as the demonstrator in a previous tool use study (Miller et al., 2016) and thus has received a more extensive exposure to the object-dropping apparatus than the other individuals we tested.

**Figure 4:**
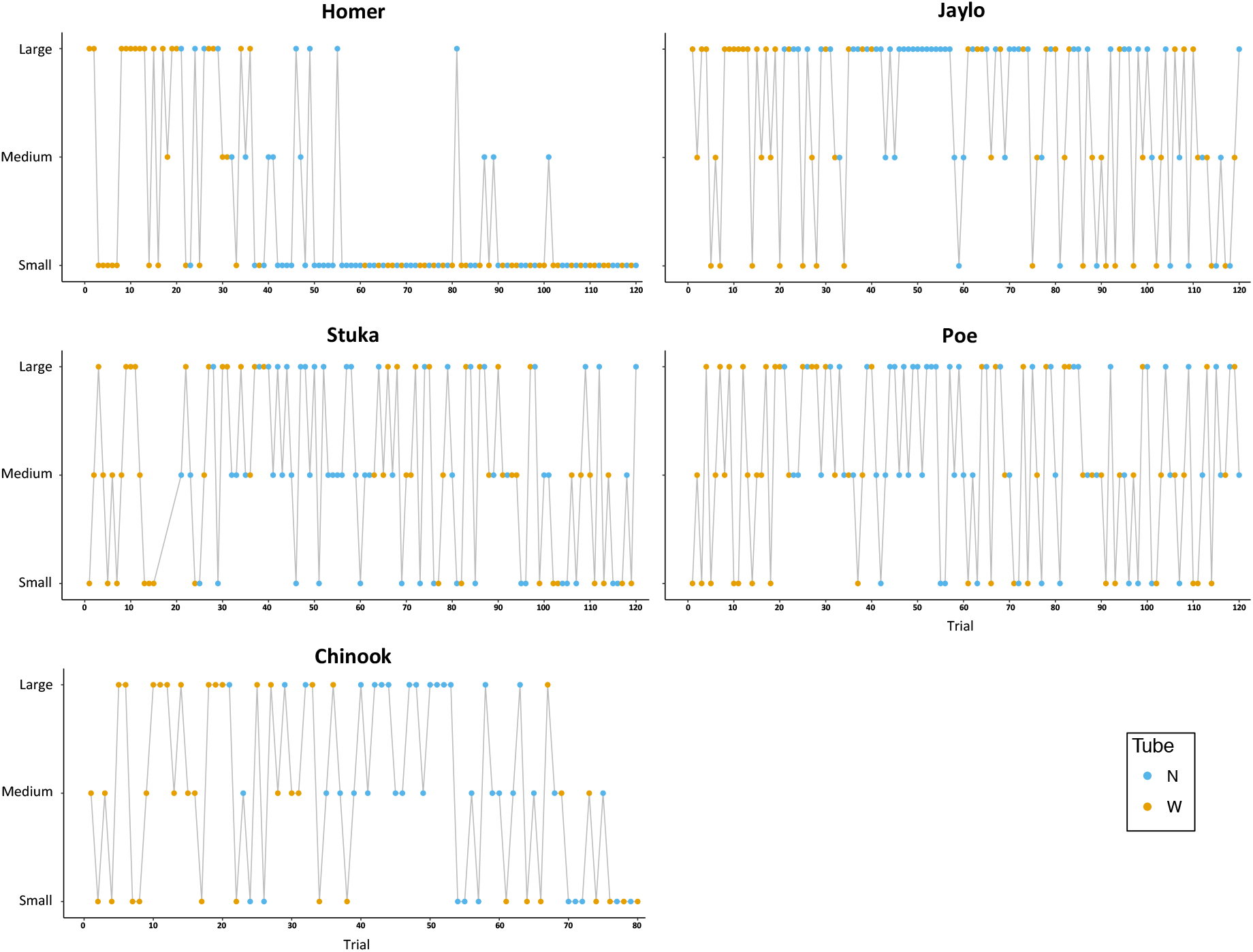
Trial by trial description of the behaviour in Experiment 1a (Block 1-4) and 1b (Block 5, 6). The apparatus that was used in each trial is also showed (wide tube: yellow dots; narrow tube: blue dots).

**Figure 5:**
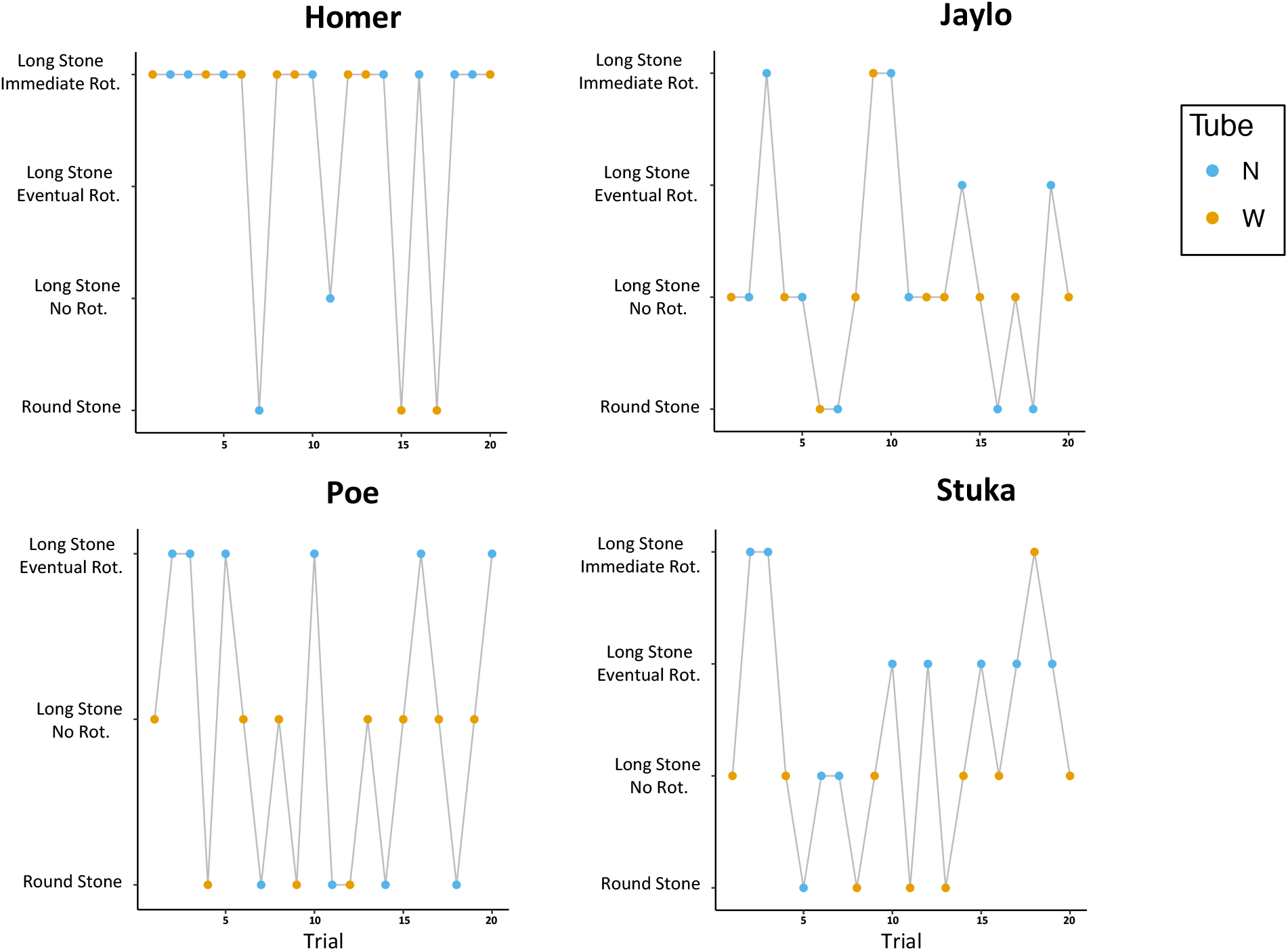
Trial by trial description of the behaviour in Experiment 2. The apparatus that was used in each trial is also showed (wide tube: yellow dots; narrow tube: blue dots).

Given that neither Eurasian jays nor rooks habitually use tools in the wild, it is interesting to compare the performance of the jays in this study with that of the rooks tested by Bird and Emery (2009). Importantly, however, one must exhibit caution in doing so given that these were two separate experiments, but tentative comparisons might yield fruit for further work in which the two groups of birds could be directly compared. When tested in a similar size selectivity test, rooks immediately switched their preference from the large stone to the small stone when the latter was the only functional tool (Bird and Emery, 2009). In the shape selectivity test, rooks, like jays in this study, expressed a pronounced preference for the long stone regardless of the condition, and higher frequencies of rotation in presence of the narrow tube (Bird and Emery, 2009). Crucially however, rooks often performed *Immediate Rotation* rather than *Eventual Rotation* in the Narrow tube condition. Taken together, the performances of these species may indicate that Eurasian jays may have more limited tool selectivity abilities than rooks (Bird and Emery, 2009). However, this possibility should be considered with caution given the methodological and ontogenetic differences between our experiments and those conducted by Bird and Emery (2009). Specifically, in Bird and Emery’s (2009) selectivity tests, rooks were systematically presented with the wide tube apparatus in the first half of trials, and subsequently with the narrow tube apparatus in the remaining trials. In contrast, the jays tested in this study did not experience such a clear sequence of exposure to the two apparatuses, as the presentation of the two apparatuses most often co-occurred within the same block of trials. Although apparently minor, it cannot be excluded that this methodological difference may have influenced the performances of the two species. Another potentially relevant difference between the studies is that jays tested in this study were juveniles (1.5 years old at test date) whereas the rooks tested by Bird and Emery (2009) were adults at the time of testing.

In regard to the stick tool test jays’ performance appears to be similar to that reported in rooks (Bird and Emery, 2009). Both species appear to exhibit good levels of flexibility in using novel tools to solve a familiar task. Supporting previous findings in rooks (Bird and Emery, 2009), jays’ use of sticks as tools also indicate that New Caledonian crow-like adaptations in beak morphology and vision (Martinho et al., 2014; Matsui et al., 2016; Troscianko et al., 2012) are not essential to achieve basic manipulations of stick tools.

Ultimately, to directly assess the question of whether the sophisticated physical cognition reported in the *Corvus* genus is shared with more distantly related species of corvids, future work will have to encompass large-scale comparative studies, where different species can be directly compared using the same or equivalent methodology.

In summary, after being trained to use stones as tools, the Eurasian jays were able to generalise to using sticks and to adopt a novel technique on the same apparatus, i.e. collapsing the internal platform by actively pushing a tool against it. What appears to be in contrast to the previously reported results for rooks and New Caledonian crows is that the Eurasian jays failed to immediately adjust their selection of tools according to the their functionality. However, the jays’ performance indicates that these birds were capable of learning to optimise their behaviour, as they progressively developed a preference for the smaller size tool, which was the only tool that was functional in both conditions, and performed the required manipulation of the functional long-shaped tool.

## Authors Contribution

SJ and PA designed the study with the help of NSC. PA collected and analysed the data. The first draft of the manuscript was written by PA with critical additions and revisions by LO, NSC and SJ. MB provided essential comments during revisions of the manuscript.

## Acknowledgments

P.A. received financial support by the Accademia Nazionale dei Lincei. S.A.J., M.B. and N.S.C. were funded by the European Research Council under the European Union’s Seventh Framework Programme (FP7/2007-2013)/ERC Grant Agreement No. 3399933, awarded to N.S.C.

## Conflict of Interest

None declared.

## Notes

https://doi.org/10.5281/zenodo.3471706

## References

Amodio, P., Jelbert, S. A., and Clayton, N. S. (2018). The interplay between psychological predispositions and skill learning in the evolution of tool use. Curr. Opin. Behav. Sci. 20, 130–137. doi:10.1016/j.cobeha.2018.01.002.

Auersperg, A. M. I., von Bayern, A. M. P., Gajdon, G. K., Huber, L., and Kacelnik, A. (2011). Flexibility in problem solving and tool use of kea and new caledonian crows in a multi access box paradigm. PLoS One 6. doi:10.1371/journal.pone.0020231.

Bird, C. D., and Emery, N. J. (2009). Insightful problem solving and creative tool modification by captive nontool-using rooks. Proc. Natl. Acad. Sci. U. S. A. 106, 10370–10375. doi:10.1073/pnas.0901008106.

Chappell, J., and Kacelnik, A. (2002). Tool selectivity in a non-primate, the New Caledonian crow (Corvus moneduloides). Anim. Cogn. 5, 71–78. doi:10.1007/s10071-002-0130-2.

Cheke, L. G., Bird, C. D., and Clayton, N. S. (2011). Tool-use and instrumental learning in the Eurasian jay (Garrulus glandarius). Anim. Cogn. 14, 441–455. doi:10.1007/s10071-011-0379-4.

Emery, N. J., and Clayton, N. S. (2004). The mentality of crows: convergent evolution of intelligence in corvids and apes. Science 306, 1903–1907. doi:10.1126/science.1098410.

Haubo, R., and Christensen, B. (2018). Ordinal: Regression Models for Ordinal Data. http://www.cran.r-project.org/package=ordinal.

Holzhaider, J., Gray, R., and Hunt, G. (2010). The development of pandanus tool manufacture in wild New Caledonian crows. Behaviour 147, 553–586. doi:10.1163/000579510X12629536366284.

Jelbert, S. A., Hosking, R. J., Taylor, A. H., and Gray, R. D. (2018). Mental template matching is a potential cultural transmission mechanism for New Caledonian crow tool manufacturing traditions. Sci. Rep. 8, 8956. doi:10.1038/s41598-018-27405-1.

Jelbert, S. A., Miller, R., Schiestl, M., Boeckle, M., Cheke, L. G., Gray, R. D., Taylor, A. H. & Clayton, N. S. (2019). New Caledonian crows infer the weight of objects from observing their movements in a breeze. Proc. R. Soc. B Biol. Sci. 286, 20182332. doi:10.1098/rspb.2018.2332.

Kabadayi, C., and Osvath, M. (2017). Ravens parallel great apes in flexible planning for tool-use and bartering. Science 357, 202–204. doi:10.1126/science.aam8138.

Kenward, B., Rutz, C., Weir, A. A. S., and Kacelnik, A. (2006). Development of tool use in New Caledonian crows: inherited action patterns and social influences. Anim. Behav. 72, 1329–1343. doi:10.1016/j.anbehav.2006.04.007.

Kenward, B., Weir, A. A. S., Rutz, C., and Kacelnik, A. (2005). Tool manufacture by naive juvenile crows. Nature 433, 121. doi:10.1038/nature03294.

Klump, B. C., Cantat, M., and Rutz, C. (2019). Raw-material selectivity in hook-tool-crafting New Caledonian crows. Biol. Lett. 15, 20180836. doi:10.1098/rsbl.2018.0836.

Klump, B. C., Masuda, B. M., St Clair, J. J. H., and Rutz, C. (2018). Preliminary observations of tool-processing behaviour in Hawaiian crows Corvus hawaiiensis. Commun. Integr. Biol. 11, e1509637. doi:10.1080/19420889.2018.1509637.

Knaebe, B., Taylor, A. H. G., Elliffe, D. M., and Gray, R. D. (2017). New Caledonian crows show behavioural flexibility when manufacturing their tools. Behaviour 154, 65–91. doi:10.1163/1568539X-00003411.

Logan, C. J., Harvey, B. D., Schlinger, B. A., and Rensel, M. (2016). Western scrub-jays do not appear to attend to functionality in Aesop’s Fable experiments. PeerJ 4, e1707. doi:10.7717/peerj.1707.

Martinho, A., Burns, Z. T., Von Bayern, A. M. P., and Kacelnik, A. (2014). Monocular tool control, eye dominance, and laterality in new caledonian crows. Curr. Biol. 24, 2930–2934. doi:10.1016/j.cub.2014.10.035.

Matsui, H., Hunt, G. R., Oberhofer, K., Ogihara, N., McGowan, K. J., Mithraratne, K., et al. (2016). Adaptive bill morphology for enhanced tool manipulation in New Caledonian crows. Sci. Rep. 6, 22776. doi:10.1038/srep22776.

Miller, R., Logan, C. J., Lister, K., and Clayton, N. S. (2016). Eurasian jays do not copy the choices of conspecifics, but they do show evidence of stimulus enhancement. PeerJ 4, e2746. doi:10.7717/peerj.2746.

R Core Team and contributors worldwide (2018). The R Stats Package. http://cran.r-project.org/web/packages/STAT/index.

RStudio Team (2018). RStudio: Integrated development environment for R. http://www.rstudio.org.

Rutz, C. (2016). Discovery of species-wide tool use in the Hawaiian crow. Nature 537. doi:10.1038/nature19103.

Rutz, C., and St Clair, J. J. H. (2012). The evolutionary origins and ecological context of tool use in New Caledonian crows. Behav. Processes 89, 153–165. doi:10.1016/j.beproc.2011.11.005.

Shettleworth, S. J. (2010). Cognition, evolution, and behavior. Oxford University Press.

Taylor, A. H., Hunt, G. R., Holzhaider, J. C., and Gray, R. D. (2007). Spontaneous Metatool Use by New Caledonian Crows. Curr. Biol. 17, 1504–1507. doi:10.1016/J.CUB.2007.07.057.

Troscianko, J., von Bayern, A. M. P., Chappell, J., Rutz, C., and Martin, G. R. (2012). Extreme binocular vision and a straight bill facilitate tool use in New Caledonian crows. Nat. Commun. 3, 1110. doi:10.1038/ncomms2111.

von Bayern, A. M. P., Danel, S., Auersperg, A. M. I., Mioduszewska, B., and Kacelnik, A. (2018). Compound tool construction by New Caledonian crows. Sci. Rep. 8, 15676. doi:10.1038/s41598-018-33458-z.

Wimpenny, J. H., Weir, A. A. S., Clayton, L., Rutz, C., and Kacelnik, A. (2009). Cognitive Processes Associated with Sequential Tool Use in New Caledonian Crows. PLoS One 4, e6471. doi:10.1371/journal.pone.0006471.

